# Oxygen-glucose deprivation and reoxygenation on human cerebral organoids alters expression related to lipid metabolism

**DOI:** 10.1101/2020.03.31.017335

**Authors:** Naoki Iwasa, Takeshi K. Matsui, Naritaka Morikawa, Yoshihiko M. Sakaguchi, Tomo Shiota, Naohiko Iguchi, Yuhei Nishimura, Eiichiro Mori, Kazuma Sugie

## Abstract

Ischemic stroke is one of the most common neurological disease. However, the impact of ischemic stroke on human cerebral tissue remains largely unknown; due to a lack of ischemic human brain samples. In this study, we used cerebral organoids derived from human induced pluripotent stem cells to evaluate the effect of oxygen-glucose deprivation/reoxygenation (OGD/R). We identified 15 differentially expressed genes (DEGs); and found that all the DEGs were downregulated. Pathway analysis showed the relationship of vitamin digestion and absorption, fat digestion and absorption, peroxisome proliferator-activated receptor signaling pathway, and complement and coagulation cascades. These findings indicate the mechanisms underlying ischemic injury in human cerebral tissue.

## Introduction

Stroke, a cerebrovascular disease, is a common neurological disorder, and ischemic strokes are along the most major causes of permanent morbidity and disability. (Spescha et al., 2013;Knowland et al., 2014;Benjamin et al., 2019). Several therapeutic agents are available for acute ischemic stroke, including thrombolytics, anticoagulants and antiplatelets. However, their efficacy is restricted and their use is limited (Catanese et al., 2017). Mechanical thrombolytics and recombinant tissue plasminogen activator (rt-PA) have therapeutic time windows for hyperacute ischemic stroke, because the target of these therapies is to rescue penumbra around ischemic core (Fisher and Bastan, 2012;Dorado et al., 2014). Thus, elucidating the mechanism of ischemic stroke in human cerebral tissue is important for the development of appropriate therapies.

Thus far, two-dimensional neuron cultures have been utilized as ischemic models. However, there are limitations to these models as actual brain tissues are composed of multiple cell types and the response to ischemia takes place from cell to cell. Recent progress in the development of organs in--a-dish (organoids) provides potential for the modeling of various diseases (Clevers, 2016). Organoids resemble the structure of organs that are composed of various cells, and three dimensionally cultured cerebral organoids are expected to represent physiological environment (Pasca et al., 2015). In this study, we analyzed the gene expression profiles of human cerebral organoids after oxygen-glucose deprivation/reoxygenation (OGD/R), an ischemic model *in vitro* (Hossmann, 1998).

## Methods

### Cell culture

iPS cells were cultivated in mTeSRTM1 medium (catalog # 05851, Stemcell Technologies, Vancouver, British Columbia, Canada) and maintained in feeder-free condition with mTESR-TM1 media. Feeder-free human iPSC line (XY) from Takara was obtained on six well plates (catalog # 3506, Corning, New York, USA) coated with growth factors reduced Matrigel (catalog # 356230, BD Biosciences, San Jose, CA, USA). At the time of passage, we added ROCK inhibitor (final concentration 10 µM, catalog # S-1049, Selleck Chemicals, Houston, Texas, USA), and maintained these cells with daily medium change without ROCK inhibitor until they reached about 70% confluency. Then, they were detached by versene solution (catalog # 15040-066, Thermo Fisher Scientific, Waltham, MA, USA) and seeded by 1:20 dilution ratio.

### Cerebral organoids generation

Cerebral organoids were differentiated according to previously published protocol (Lancaster et al., 2013). Human iPSCs were detached and subjected to embryoid body (EB) induction using the protocol. After four days, half of the media was replaced by human EB medium without ROCK inhibitor and basic FGF. Two days later, the EBs were transferred into a neural induction media, and then embedded in Matrigel after five days. Subsequently, the organoids were induced in organoid medium by using an orbital shaker. The total process took 28days, from EB formation.

### Immunohistochemical analysis

Human cerebral organoids were fixed in 4% paraformaldehyde in Phosphate-Buffered Saline (PBS) overnight at 4°C, dehydrated with 30% sucrose in PBS and embedded in OCT Compound (23-730-571, Thermo Fisher Scientific). Cryostat sections (14 μm) were cut and mounted on slides (Thermo Fisher Scientific). Mounted sections were incubated at room temperature for 1 hour with blocking solution (3% normal donkey serum+0.3% Triton X-100 in Tris-Buffered Saline [TBS]) and subsequently incubated with primary antibodies diluted in blocking solution overnight at 4 °C. After three washes with TBS, corresponding fluorophore-conjugated secondary antibodies diluted in the blocking solution were added and samples were incubated at room temperature for 2 hours. Finally, stained slides were first rinsed three times with TBS, then mounted and analyzed using a microscope. Antibodies specific for TUJ1 (1:400, T8660, Sigma) and SOX2 (1:400, 23064S, Signaling) were used for immunostaining.

### Oxygen-glucose deprivation/reoxygenation

The organoids were plated in DMEM (09891-25, NAKARAI) without glucose, and 1% (v/v) penicillin-streptomycin. Subsequently, the organoids were cultured in a hypoxic incubator (94% N_2_, 5% CO_2_, 1% O_2_) at 37 °C for 1 hour. For reoxygenation, the medium was changed to Primary Neuron Basal Medium, then incubated (95% air, 5% CO_2_) at 37 °C for 1hour.

### RNA isolation

RNA from human cerebral organoids was extracted according to the manufacturer’s protocol supplied with TRIzol reagent (15596018, Thermo Fisher Scientific). The concentration and purity of the RNA samples were measured using Spectrophotometer (Beckman Coulter).

### RNA sequencing

Total RNA was isolated from the cells using the PureLink RNA Mini Kit (12183018A) and according to the manufacturer’s instructions. RNA concentration was analyzed by Qubit RNA HS Assay Kit (Thermo Fisher Scientific) and the purity was assessed using the Qsep100 DNA Fragment Analyzer and RNA R1 Cartridge (BiOptic). Then, total RNA was converted to cDNA and used for Illumina sequencing library preparation, according to the KAPA Stranded mRNA-Seq Kit protocols (KAPA BIOSYSTEMS). DNA fragments were ligated by FastGene Adapter Kit (FastGene). After the purified cDNA library products were appreciated using Qubit dsDNA HS Assay Kit (Thermo Fisher Scientific), they were qualitatively evaluated using the Fragment Analyzer and dsDNA 915 Reagent Kit (Advanced Analytical Technologies), before it was finally sequenced (2×76bp) on NextSeq 500 (Illumina). Sickle (ver.1.33) is an error correction tool for quality trimming. It discards reads based upon the length threshold derived from quality score for FASTQ files. Using Sickle (ver.1.33), the threshold identified were quality values under 20base and reads under 30 bp. Additionally, Hisat2 software was run with default parameters. The sequence alignment files generated by Sickle and Hisat2 were used to generate counts of mapped reads.

### RT-PCR and quantitative PCR

Extracted RNA samples were either sent to Bioengineering lab for RNA-sequencing analysis, and or subjected to RT-PCR. For RT-PCR, the extracted RNAs were reverse transcribed using the protocol supplied with ReverTra Ace qPCR RT Master Mix (FSQ-201, TOYOBO). StepOne Plus Real-time PCR System (Thermo Fisher Scientific) was used to amplify and quantify levels of target gene cDNA. Quantitative real-time PCR (qRT-PCR) was conducted using SsoAdvanced Universal SYBR Green Supermix (172-5271, Bio-Rad Laboratories) and specific primers for qRT-PCR.

Primers used in this study are:;

FTL [F: 5′-TACGAGCGTCTCCTGAAGATGC-3′;

R: 5′-GGTTCAGCTTTTTCTCCAGGGC-3′],

AHSG [F: 5′-ACGCTCAGAACAACGGCTCCAA-3′;

R: 5 ′-GCAACACAGTCAGTGCCAGACA-3′],

FGG [F: 5′-GAAGGCAACTGTGCTGAACAGG-3′;

R: 5′-CCATTAGGAGTAGATGCTTTTGAG-3′],

TTR [F: 5′-CGTGCATGTGTTCAGAAAGGCTG-3′;

R: 5′-CTCCTCAGTTGTGAGCCCATGC-3′], and

AFP [F: 5′-GCAGAGGAGATGTGCTGGATTG-3′;

R: 5′-CGTGGTCAGTTTGCAGCATTCTG-3′].

Reactions were run in triplicate. The expression of each gene was normalized to the geometric mean of β-actin as a housekeeping gene and analyzed using the ΔΔCT method.

### Differentially expressed genes (DEGs) extraction

By comparing organoids under OGD/R condition to non-treated organoids and using R package edgeR (Robinson et al., 2010) in R software (version 3.5.3), DEGs were extracted. The MA plot in R is a visual tool for showing the total gene expression levels of DEGs. The X-axis represents log CPM – log counts per million – and are measures of gene expression level. The Y-axis indicates log FC. Log FC is the log fold-change, which in this case, is the log difference between cerebral organoids after OGD/R.

### Gene ontology and pathway analysis

Using the Database for Annotation, Visualization, and Integrated Discovery (DAVID) version 6.8 database (https://david.ncifcrf.gov/), selected DEGs were analyzed. The DAVID database shows the molecular function, biological process, and cellular component expressed in the gene profile. In this study, the DAVID database was applied to investigate gene ontological (GO) annotation and Kyoto encyclopedia of genes and genomes (KEGG) pathways of DEGs. P-value <0.05 was chosen as the threshold of KEGG pathway. Upregulation and downregulation of genes were analyzed separately.

### Protein-Protein Interaction (PPI) analysis

Based on the Search Tool for the Retrieval of Interacting Genes (STRING, https://string-db.org/; Szklarczyk et al., 2019). Online database, DEGs were inputted and the PPI network was constructed. The STRING database shows the interaction of each protein from prediction or experiments. The threshold to construct the PPI network was confidence score >0.4.

### Statistical analysis

Statistical analysis was performed using the GraphPad Prism7 (GraphPad Software). A one-way analysis of variance (ANOVA) was applied to the q-PCR data was analyzed one-way analysis of variant (ANOVA). P-value <0.05 was defined as the threshold.

## Results

To generate a model of cerebral ischemia/reperfusion *in vitro*, cerebral organoids were treated with oxygen-glucose deprivation/reoxygenation (OGD/R), which is a well-established method of mimicking the pathological processes of ischemia. We first cultured the human cerebral organoids for 28 days. We then validated that fluorescent micrographs showed a coexistence of TUJ1 (green), SOX2 (red) and DAPI (blue) in human cerebral organoids (Figure 1).

**Figure 1.**
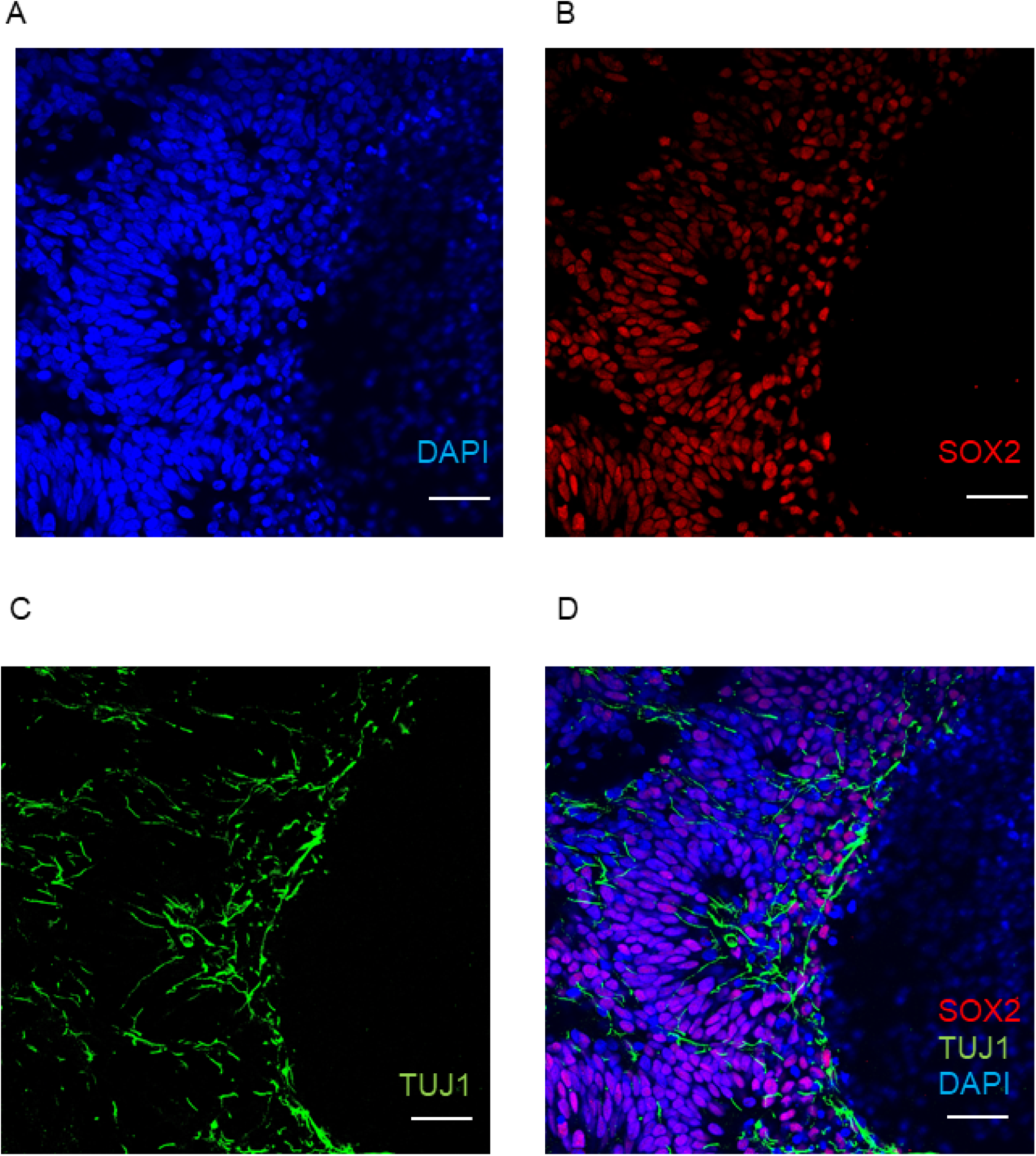
Imunohistochemical staining of non-treated human cerebral organoids. Immunohistochemical staining of neuronal cells’ marker (TUJ1), embryonic stem cells’ marker (SOX2) and 49,6-diamidino-2phenylindole (DAPI). A Immunohistochemical staining by DAPI B Immunohistochemical staining by SOX2 C Immunohistochemical staining by TUJ1 D Immunohistochemical costaining by DAPI, SOX2 and TUJ1 Fluorescent micrographs show coexisting of TUJ1 (green), SOX2 (red) and DAPI (blue) at human cerebral organoids. Bars = 100µm.

Subsequently, these organoids were placed under OGD condition for 1 hour and reoxygenation for 1 hour. After OGD/R, RNA was extracted (Figure 2A) prior to RNA isolation and RNA sequencing, whereupon gene ontology analysis was performed. Significant DEGs were colored in red (Figure 2B). Using the threshold of p<0.05 and |log FC| >0.1, a total of 52 DEGs were determined between intact cerebral organoids and cerebral organoids under OGD/R, including 14 upregulated and 38 downregulated genes (Table 1). When false discovery rate (FDR) adjusted p-value ≤ 0.05 was used as a threshold, 15 genes of AFP, TTR, APOA2, ALB, APOA1, RNA28SN4, APOC3, FTL, AHSG, FGG, FABP1, MIR3615, FGB, RNA45SN4 and FGA were detected and all genes were found to be downregulated (Table 1). The top 5 significant downregulated genes were AFP, TTR, APOA2, ALB and APOA1 (Table 1). Though the expression level of these five genes were also confirmed by quantitative PCR (q-PCR), they were not significant (P > 0.05) (Figure 3).

**Table 1.**
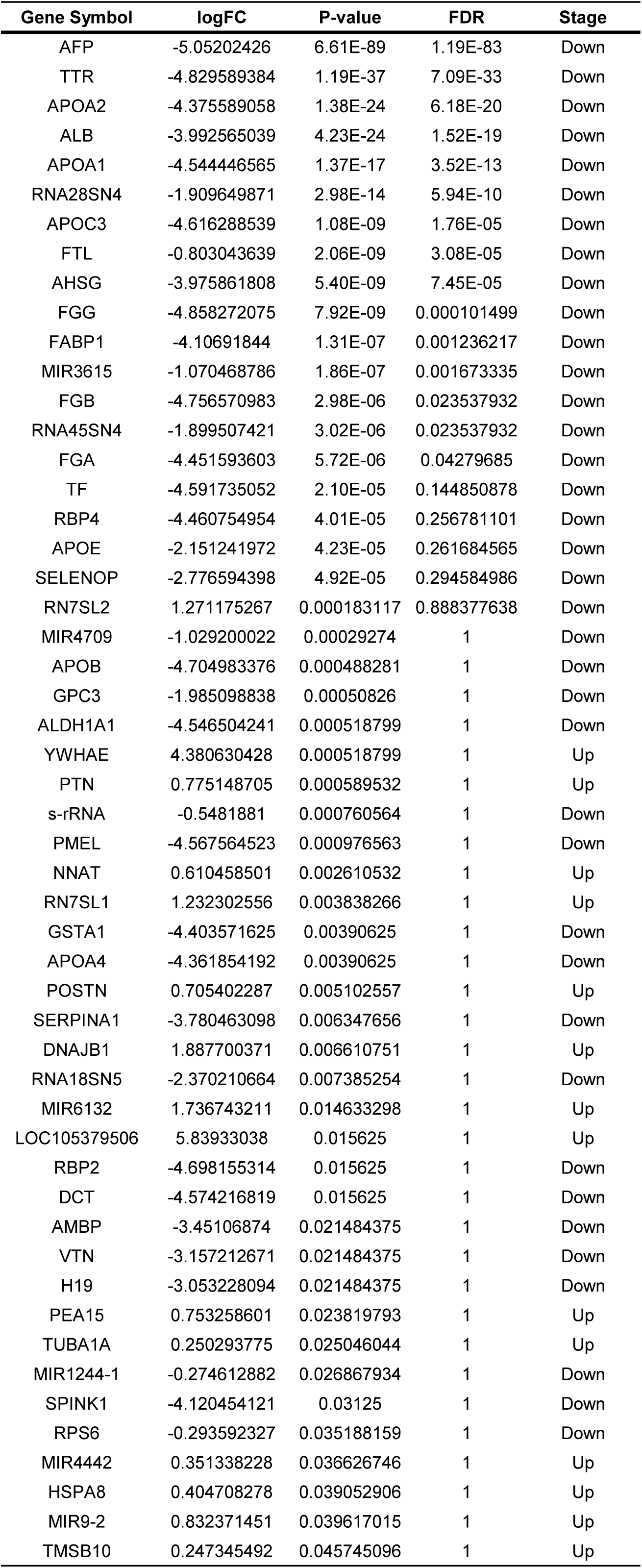
Profiling gene expression comparing organoids under OGD/R condition with non-treated organoids

**Figure 2.**
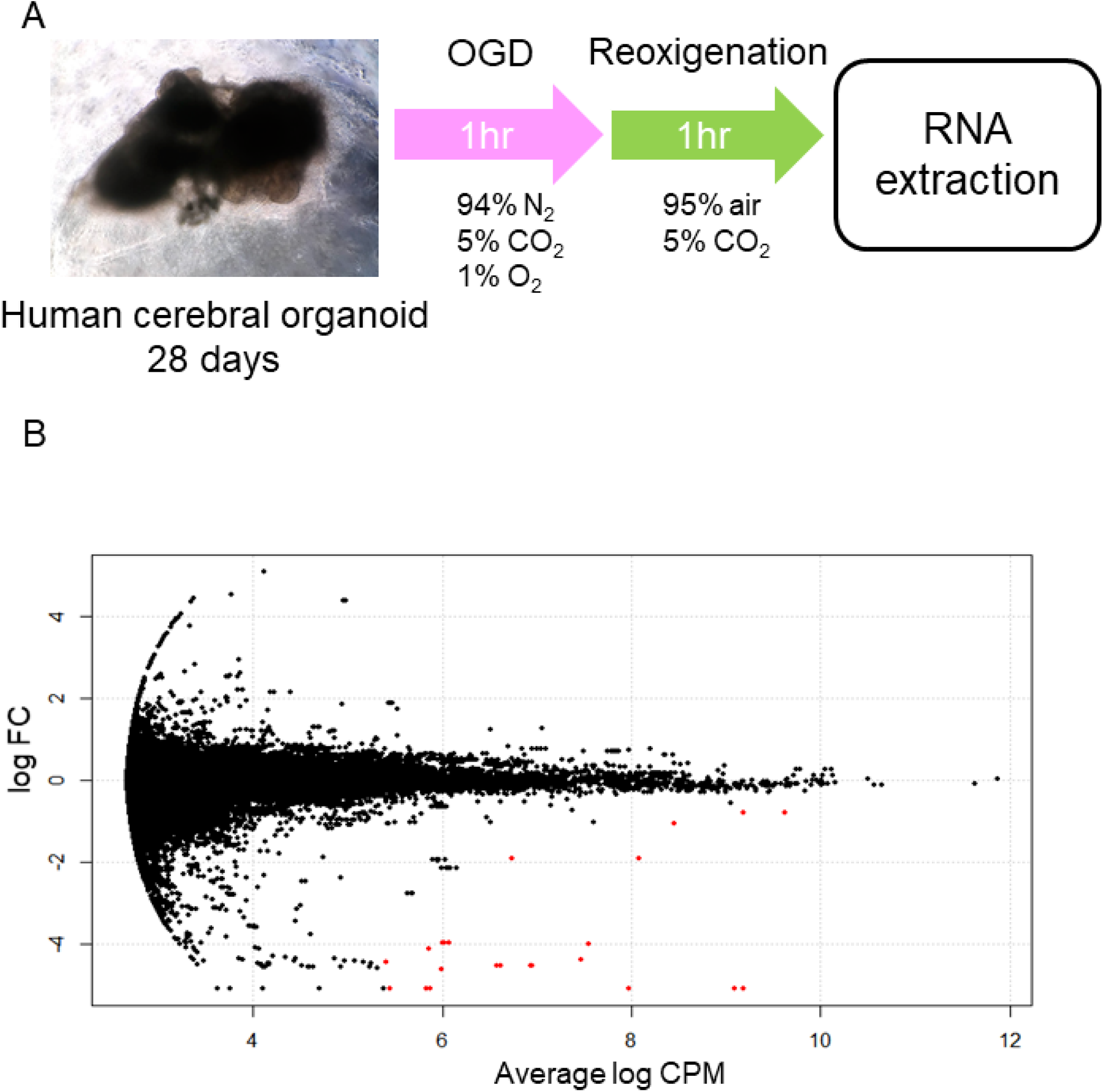
**A. Schema of this experimental procedure.** First, human cerebral organoids were generated and cultured for 28 days. Subsequently, the organoids were cultured in a hypoxic incubator (94% N_2_, 5% CO_2_, 1% O_2_) at 37 °C for 1 hour. For reoxygenation, they were incubated (95% air, 5% CO_2_) at 37 °C for 1 hour. After OGD/R treatment, RNA was extracted. **B. MA plot of cerebral organoids.** The X-axis is log CPM, which is a measure of gene expression level. The Y-axis indicate log FC, which is the log difference between cerebral organoids after OGD/R and controls. Red plots are significant DEGs. CPM: counts per million, FC: fold-change, DEGs: differentially expressed genes

**Figure 3.**
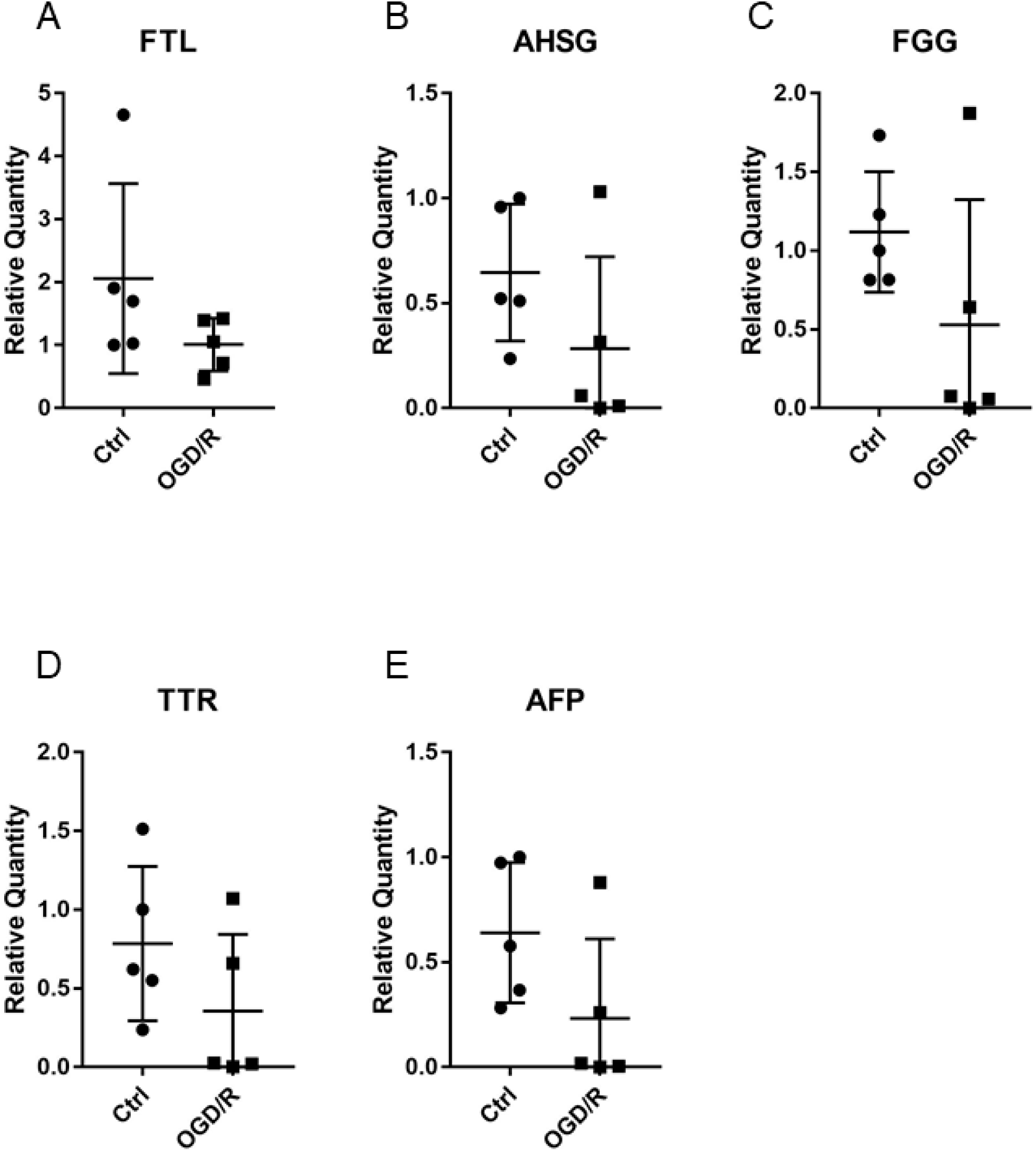
qPCR analysis of cerebral organoids [control VS OGD/R condition]. Gene expression analysis of FTL (A), AHSG(B), FGG(C), TTR(D), and AFP(E) on non-treated organoids (Ctrl) VS organoids after OGD/R (OGD/R). The P-values were 0.0862 (FTL), 0.0881 (AHSG), 0.0869 (FGG), 0.103 (TTR) and 0.0554 (AFP).

Gene ontological (GO) analysis in the biological process category showed that OGD/R on human cerebral organoids induced lipid metabolism and blood coagulation (Table 2). The pathway-based analysis, Kyoto encyclopedia of genes and genomes (KEGG) pathway, further highlighted the vitamin digestion and absorption, fat digestion and absorption, peroxisome proliferator-activated receptor (PPAR) signaling pathway, and complement and coagulation cascades (Table 3). Using the Search Tool for the Retrieval of Interacting Genes (STRING) online database, 52 DEGs were filtered into the protein-protein interaction (PPI) network of DEGs containing nodes and edges with parameters including a minimum required interaction score>0.4 (medium confidence) (Figure 4).

**Table 2.** Go term analysis derived from biological process of human cerebral organoids after OGD/R (FDR<0.05)

| Term | Count | P-value | FDR |
| --- | --- | --- | --- |
| retinoid metabolic process | 11 | 5.91E-17 | 1.55E-13 |
| platelet degranulation | 9 | 5.07E-11 | 7.27E-08 |
| lipoprotein metabolic process | 7 | 1.96E-10 | 2.81E-07 |
| cholesterol efflux | 6 | 2.02E-09 | 2.89E-06 |
| lipoprotein biosynthetic process | 5 | 2.49E-09 | 3.58E-06 |
| phospholipid efflux | 5 | 1.97E-08 | 2.82E-05 |
| high-density lipoprotein particle remodeling | 5 | 2.68E-08 | 3.84E-05 |
| reverse cholesterol transport | 5 | 5.97E-08 | 8.56E-05 |
| triglyceride catabolic process | 5 | 2.44E-07 | 3.50E-04 |
| cholesterol homeostasis | 6 | 2.72E-07 | 3.90E-04 |
| high-density lipoprotein particle assembly | 4 | 5.47E-07 | 7.85E-04 |
| positive regulation of cholesterol esterification | 4 | 8.20E-07 | 0.0011757 |
| protein polymerization | 4 | 2.77E-06 | 0.0039787 |
| negative regulation of very-low-density lipoprotein particle remodeling | 3 | 1.42E-05 | 0.0202973 |
| cholesterol metabolic process | 5 | 1.47E-05 | 0.0210621 |
| lipid transport | 5 | 2.28E-05 | 0.0327645 |
| triglyceride homeostasis | 4 | 2.47E-05 | 0.0354574 |
| blood coagulation, fibrin clot formation | 3 | 2.83E-05 | 0.0405344 |

**Table 3.** KEGG pathway of differentially expressed genes.

| **Term** | **Count** | **Genes** | **FDR** | **P-value** |
| --- | --- | --- | --- | --- |
| Vitamin digestion and absorption | 4 | APOA4, APOB, APOA1, RBP2 | 0.058366266 | 6.24E-05 |
| Fat digestion and absorption | 4 | APOA4, APOB, APOA1, FABP1 | 0.332117918 | 3.56E-04 |
| PPAR signaling pathway | 4 | APOA2, APOA1, APOC3, FABP1 | 1.618503332 | 0.00174259 |
| Complement and coagulation cascades | 4 | FGG, FGA, FGB, SERPINA1 | 1.760575716 | 0.00189677 |
| Platelet activation | 3 | FGG, FGA, FGB | 54.24273463 | 0.08016934 |

**Figure 4.**
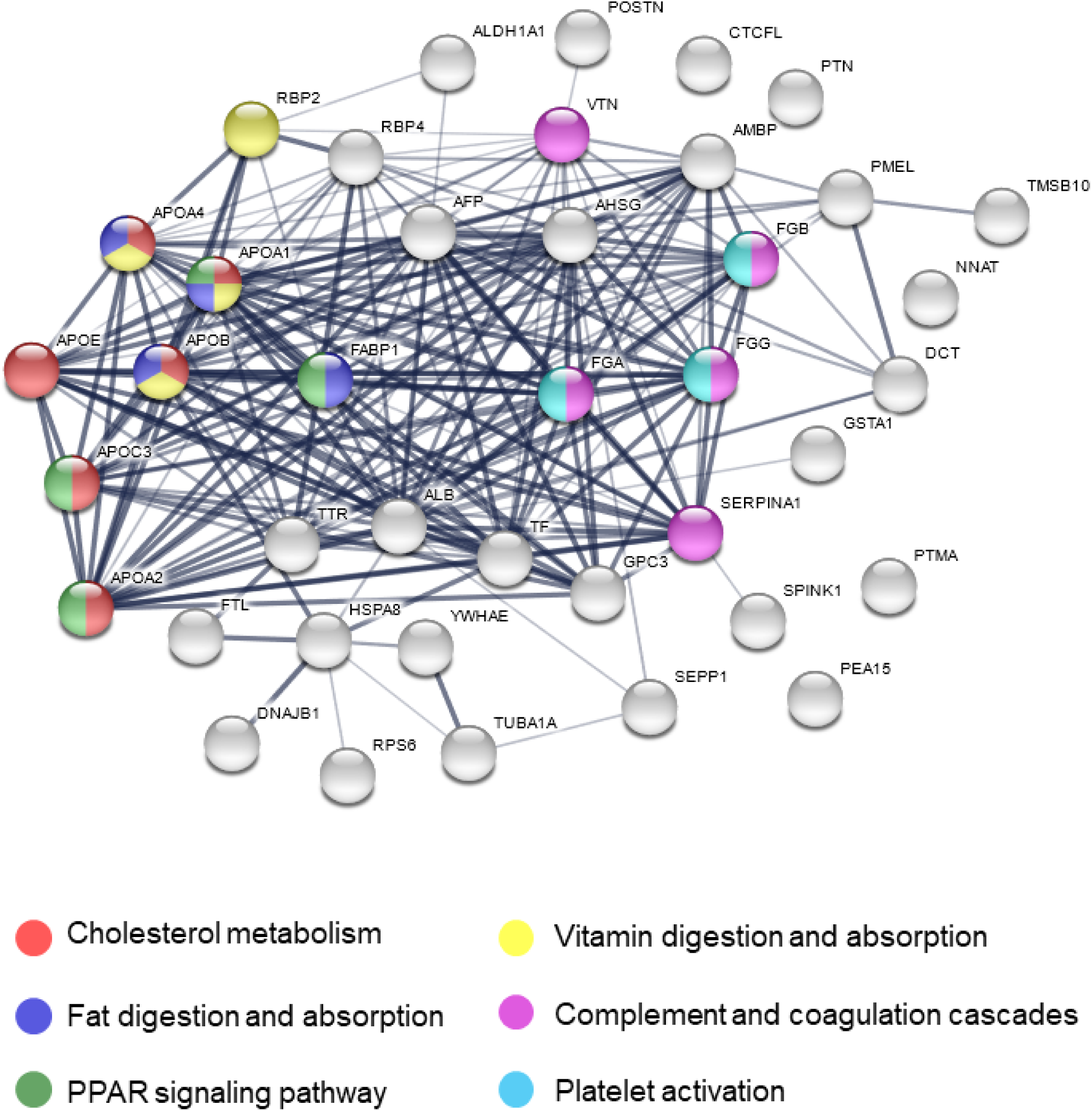
Protein-Protein Interaction network of differentially expressed genes. Nodes are proteins and the thickness of lines indicates strong interaction between proteins. Colored nodes mean related pathway.

## Discussion

In rat models of acute pneumococcal meningitis, genes related to neurotransmission and lipid metabolite are downregulated (Coimbra et al., 2006). Bacterial meningitis leads to sugar reduction in cerebral spinal fluid (CSF) and cerebral vasculitis, thus mimicking ischemia. In this study, 15 genes and several pathways were identified using analysis of DEGs (Table 1). Many genes associated with lipid metabolism were observed to be downregulated, APOA1, APOA2, APOC3, and FABP1 (Table 1). PPAR-α agonists work protectively against ischemia or trauma (Besson et al., 2005; Collino et al., 2006). PPAR-γ agonists conduct neuroprotection against focal ischemia in the rat brain (Zhao et al., 2005). Dual PPARα/γ agonist improves stroke outcome after transient cerebral ischemia in mice (Boujon et al., 2019). These results suggest that therapeutic agents related lipid metabolism, including PPAR agonists, may be effective for ischemia.

Vascular endothelial growth factor (VEGF) and S100A12 are elevated in patients with acute ischemic stroke (Stowe et al., 2007; Li et al., 2014; Wakisaka et al., 2014; Billinger et al., 2018). VEGF increases are especially acute in peri-infarct region following cortical infarct (Stowe et al., 2007). In our study, we also confirmed the VEGF and S100A12 elevation after OGD/R condition. In addition, we detected a number of upregulated genes—RN7SL2, YWHAE, PTN, NNAT, RN7SL1, POSTN, DNAJB1, MIR6132, LOC105379506 and PEA15 (Table 1). In the brain, YWHAE is involved in directing the movement of nerve cells by binding to other proteins (Toyo-oka et al., 2003). Pleiotrophin (PTN) plays important roles in cell growth and survival, cell migration, and the expression of inflammatory cytokines (Shen et al., 2017). YWHAE and PTN are mainly expressed in the cerebral cortex (Kido et al., 2014; Shen et al., 2017). POSTN leads to decreased apoptosis during hypoxia (Aukkarasongsup et al., 2013). Results from these studies indicate that upregulated genes might be associated with neuroprotection.

Reduction of albumin and AFP were associated with OGD/R (P-value<0.05) (Table 1). The two main plasma proteins found in CSF during embryonic period is albumin and AFP (Dziegielewska et al., 1981). During the initial stages of the generation of cerebral cortex neurons, the expression of albumin and AFP were found in the outer half of cortical plate (Mollgard and Jacobsen, 1984). Normal matured human brains typically do not express albumin and AFP, and the downregulation of albumin and AFP in human cerebral organoids after OGD/R in this study was most likely due to the immaturity of the human cerebral organoids we used; we used one-month cultured immature human cerebral organoids for ischemic tissues. Nonetheless, we believe our current study paves the way to understanding the mechanism underlying ischemic stroke in human cerebral tissues. Further technical advancements will provide us a more comprehensive understanding of human brain function and disease.

## Acknowledgements

The authors thank Keren-Happuch E Fan Fen for her critical reading of the manuscript. This work was supported by grants from JSPS KAKENHI [JP17H07031 to E.M., JP19K16925 to T.K.M.], AMED The Program for Technological Innovation of Regenerative Medicine [JP19bm0704039h to T.K.M.], AMED Osaka University Seeds (A) to T.K.M, Takeda Science Foundation to E.M. and T.K.M., Kanzawa Medical Research Foundation to E.M., Uehara Memorial Foundation to E.M., Nakatomi Foundation to E.M., Konica Minolta Science and Technology Foundation to E.M., Naito Foundation to E.M., MSD Life Science Foundation to E.M., Mochida Memorial Foundation for Medical and Pharmaceutical Research to E.M., SENSHIN Medical Research Foundation to E.M., Terumo Foundation for Life Sciences and Arts to E.M., Nara Kidney Disease Research Foundation to E.M., Novartis Research Grants to E.M., K.S., Sumitomo Dainippon Pharma Research Grant to T.K.M., Nara Medical University Grant-in-Aid for Collaborative Research Projects to K.S., Nara Medical University Grant-in-Aid for Young Scientists to T.K.M., and by unrestricted funds provided to E.M. from Dr. Taichi Noda (KTX Corp., Aichi, Japan) and Dr. Yasuhiro Horii (Koseikai, Nara, Japan). The authors declare that this study received funding from parties mentioned above. The funders were not involved in the study design, collection, analysis, interpretation of data, the writing of this article or decision to submit it for publication.

## Author contributions

N.I., T.K.M, K.S., and E.M. designed the study; N.I., T.K.M, N.M., Y.M.S, N.I., K.S. and E.M. conducted the research; N.I., T.K.M, Y.M.S, Y.N., K.S. and E.M. analyzed the data; N.I., T.K.M, K.S., and E.M. wrote the paper. All authors contributed to manuscript revision, read and approved the submitted version.

